# Reduced Hippocampal-Cortical Connectivity During Memory Suppression Predicts the Ability to Forget Unwanted Memories

**DOI:** 10.1101/2022.02.08.479070

**Authors:** Yuchi Yan, Justin C. Hulbert, Kaixiang Zhuang, Wei Liu, Dongtao Wei, Jiang Qiu, Michael C. Anderson, Wenjing Yang

**Affiliations:** Key Laboratory of Cognition and Personality (SWU), Ministry of Education, Chongqing 400715, China; Faculty of Psychology, Southwest University (SWU), Chong-qing, 400715, China; Psychology Program, Bard College, Annandale-on-Hudson, New York 12504, USA; School of Psychology, Central China Normal University (CCNU), Wuhan, China; MRC Cognition and Brain Sciences Unit, University of Cambridge

**Author notes:** These authors contributed equally to this work (Michael C. Anderson, Wenjing Yang).

**Keywords:** Retrieval suppression, Inhibitory control, Functional connectivity, Hippocampus, Connectome-based predictive modeling

## Abstract

The ability to suppress unwelcome memories is important for productivity and well-being. Successful memory suppression is associated with hippocampal deactivations and a concomitant disruption of this region’s functionality. Much of the previous neuroimaging literature exploring such suppression-related hippocampal modulations has focused on the region’s negative coupling with the prefrontal cortex. In contrast, task-based changes in functional connectivity between the hippocampus and other brain regions have remained relatively underexplored. Here, we utilize psychophysiological interactions and seed connectome-based predictive modeling (seed-CPM) to investigate the relationship between the hippocampus and the rest of the brain as 134 participants attempted to suppress unwanted memories during the Think/No-Think task. The results show that during retrieval suppression, the right hippocampus exhibited decreased functional connectivity with visual cortical areas (bilateral intracalcarine cortex, right cuneal cortex, left lingual gyrus, right supracalcarine cortex, right occipital pole), left nucleus accumbens and the brain-stem that predicted superior forgetting of unwanted memories on later memory tests. Validation tests verified that prediction performance was not an artifact of head motion or prediction method and that the negative features remained consistent across different brain parcellations. These findings suggest that systemic memory suppression involves more than the modulation of hippocampal activity—it alters functional connectivity patterns between the hippocampus and visual cortex, leading to successful forgetting.

## Introduction

Some memories we relish retrieving; others risk bringing our internal life and outward productivity to a standstill if allowed into consciousness. When confronted with a reminder of a memory that threatens to cause upset, individuals may summon cognitive control processes aimed at stopping—or suppressing—retrieval of the event. With practice, such control curbs unwanted memory intrusions in the moment, as well as the likelihood of their future retrieval—outcomes linked to the modulation of hippocampal activity (Levy and Anderson, 2012; for a recent review, see Anderson and Hulbert, 2021). Much of the evidence for these mnemonic outcomes stems from a procedure known as the Think/No-Think (TNT) paradigm.

The TNT paradigm was developed to model memory suppression in the laboratory, opening an empirical window to the behavioral consequences (Anderson and Green, 2001) and neural correlates (Anderson et al., 2004) of attempts to stop memory retrieval. The paradigm consists of three main phases: an encoding phase, the critical TNT phase, and surprise memory tests. During encoding, participants are asked to learn a number of cue-target word pairs to criterion. Then, participants engage in the TNT phase, in which they repeatedly retrieve (Think condition) or suppress retrieval (No-Think condition) of the target words, when given the cue word. As reviewed in detail elsewhere (Anderson and Huddleston, 2012; Anderson and Hanslmayr, 2014; Marsh and Anderson, 2020; Anderson and Hulbert, 2021), surprise final tests for all the learned associates generally reveal that targets in the No-Think condition are less recallable than are the Baseline items, which were learned just as well at the outset, but which were omitted from the TNT phase. This below-baseline memory impairment has been termed suppression-induced forgetting (SIF).

Deployed strategically, effective memory suppression is thought to provide numerous psychological and health-related benefits (for perspectives on this topic, see Fawcett and Hulbert, 2020; Nørby, 2015), as evidenced by correlational evidence linking greater suppression abilities to reduced depression, anxiety, and PTSD symptoms (see Stramaccia et al., 2021 for a meta-analysis across these and other disorders). Whether the goal is to detect potential vulnerabilities or to train more adaptive coping habits in individuals who are or may face future challenges, understanding the nature of these relationships and their neural underpinnings is of critical concern. And just as appreciating the potential side effects of retrieval suppression can inform therapeutic approaches, their very existence has helped establish a more detailed mechanistic understanding of adaptive memory control in action (see Anderson and Hulbert, 2021, for a review).

The targets of memory suppression are not the only items that tend to be forgotten after repeated attempts to stop retrieval. Episodic memories that are encoded (Hulbert et al., 2016) or cued (Zhu and Wang, 2021) before or after periods of targeted retrieval suppression also become less accessible. This so-called “amnesic shadow” can affect memories that are entirely unrelated to the targets of retrieval suppression but simply are unlucky enough to occur near in time to suppression. This phenomenon bears a striking resemblance to organic amnesias caused by damage to the hippocampus (for reviews, see Spiers et al., 2001). In fact, the amnesic shadow was predicted based on numerous reports of hippocampal deactivations (relative to the Think condition, as well as to passive baseline conditions) observed during studies of memory suppression (Anderson et al., 2004; Depue et al., 2007; Benoit and Anderson, 2012; Paz-Alonso et al., 2013; Gagnepain et al., 2014; Benoit et al., 2016; Yang et al., 2020; Apšvalka et al., 2022). Consistent with a role of hippocampal deactivations in successful memory control, hippocampal modulation is greatest (and predictive of suppression-induced forgetting) when unwanted memories intrude and need to be purged from awareness (Levy and Anderson, 2012; Gagnepain et al., 2017).

Successful retrieval of recent memories does not depend only on hippocampal activity, however. It also depends on interactions between the hippocampus and other brain regions, such as the neocortex (Treves and Rolls, 1994; Sutherland and McNaughton, 2000; Schott et al., 2013). One prominent view of retrieval holds that its success depends on pattern completion based on perceptual inputs to the hippocampus, helps to reinstate patterns that initially encoded in the neocortex (Rolls, 2013; Horner et al., 2015; Guzman et al., 2016; Hindy et al., 2016). Indeed, the human hippocampus has extensive connections with early visual cortex, parietal cortex, and cingulate gyrus, among other regions (Huang et al., 2021). These direct connections facilitate hippocampal-cortical communication, with recall of vivid memories being associated with higher communication efficiency of the hippocampal whole-brain network (Geib et al., 2017). If suppression, rather than retrieval, is the goal, connectivity within and between memory regions might be disrupted. Neuroimaging evidence supports this prediction. For example, suppression not only reduces bilateral hippocampal activity, but also activity in posterior cortex in a content-specific manner; thus, suppression of visual objects or places reduces activity in fusiform gyrus and parahippocampal place area, respectively (Depue et al., 2007; Gagnepain et al., 2014; Benoit et al., 2015; Mary et al., 2020). Like the hippocampus, these other regions are involved in memory representation and retrieval, though for more specialized forms of content (Albers et al., 2013; Waldhauser et al., 2016; Rosenthal et al., 2016). For example, visual or auditory cortical regions involved during encoding an event are also reactivated during that event’s retrieval (Nyberg et al., 2000; Wheeler et al., 2000; Waldhauser et al., 2016). The hippocampus, as a convergence zone, integrates information from many brain areas (Backus et al., 2016), yielding increased functional connectivity between the hippocampus and the neocortex, such as sensory cortex (Ranganath et al., 2005; Wolosin et al., 2012; Schott et al., 2013). Nevertheless, most studies have focused on hippocampal connectivity changes during active recall; changes to functional connectivity linking the hippocampus to other brain areas during suppression remain relatively underexplored.

For present purposes, we examined how retrieval suppression modulates hippocampal connectivity with the rest of the brain by utilizing connectome-based predictive modeling (CPM) (Shen et al., 2017; Goldfarb et al., 2020), which previously has revealed brain functional connectivity related to attention, stress, and creative ability (Rosenberg et al., 2018; Beaty et al., 2018; Goldfarb et al., 2020). Using a cross-validation approach, CPM minimizes the chance of overfitting and improves test-retest reliability. Moreover, compared to other machine learning methods, the simplicity of the CPM approach allows for more interpretable models. Our aim was to exploit these methodological advantages to predict behavioral performance on a standard memory suppression task and to further elucidate the brain mechanisms of the associated cognitive processes.

In the current study, 134 participants completed the TNT task during functional magnetic resonance imaging (fMRI) scanning. To apply CPM, we first divided the brain into 132 regions, which combining Harvard-Oxford atlas cortical (Tzourio-Mazoyer et al., 2002), subcortical areas and cerebellar areas of automated anatomical atlas, then we selected both the right and left hippocampi as anatomical seeds. As input features for the CPM, we used a generalized psycho-physiological interaction (g-PPI) method to calculate the functional connectivity changes between seed regions with other brain regions during memory suppression (McLaren et al., 2012). Ridge regression was used to build the prediction model, allowing for different features to contribute flexibly to the prediction model.

We hypothesized that superior forgetting of unwanted memories would arise, in part, by disrupting communication between the hippocampus and posterior cortical regions involved in memory, including the visual cortex, fusiform gyrus, and the parahippocampal gyrus. Such disrupted communication should be reflected in reduced functional connectivity between the hippocampus and these regions during suppression trials. We based this hypothesis on evidence that inhibiting memory retrieval may interrupt pattern completion (Anderson et al., 2004; Depue, 2012), which, in turn, would alter hippocampal functional connectivity with regions involved in memory representation.

## Method

### Participants

We recruited 146 students from Southwest University for this study. All participants had no history of visual, medical, neurological or memory disorders and completed the TNT phase (Anderson et al., 2004) during fMRI scanning. After removing participants due to missing behavioral indices or functional scans and due to severe head motion (pre-defined as mean FD > 0.3 mm), 134 participants (Male: 41, Female: 93; average age: 19.5 years) were retained for analyses. This study has been approved by the Academic Committee of Southwest University in China. The resting-state images and behavioral data of the current 146 participants were used in another study (Yang et al., 2021). In the current study, we only considered the functional images.

### Behavioral assessment (TNT task)

Participants performed the Think/No-Think task (Anderson et al., 2004), which assesses the ability to suppress unwanted memories. The TNT task consists of three main phases: the encoding phase, the TNT phase, and the memory test phase. The TNT phase was completed in the scanner, whereas the encoding phase and memory test phases were completed outside the scanner. Further task details are provided elsewhere for space reasons (Yang et al., 2020). Here, we briefly describe the three main phases.

#### Encoding phase

During the encoding phase, participants were asked to study associations between 66 weakly related cue-target word pairs such that when presented each cue word, they would be able to reliably recall the associated target. To advance to the main TNT phase, participants were required to recall at least 50% of the target words when presented with their associated cue. Participants had up to three test-feedback cycles to demonstrate this competence after an initial study period.

#### TNT phase

After participants demonstrated successful encoding of the threshold number of cue-target word pairs, TNT practice began using filler pairs to ensure that participants understood the instructions pertaining to the Think and No-Think conditions. The training phase consisted of two blocks. After participants completed each training block, the experimenter administered a standard diagnostic questionnaire to make sure they followed the instructions correctly (Liu et al., 2021). Before entering the scanner to complete the formal TNT task, participants received a five-minute break. Then, prior to scanning, the correct word pairs were presented one final time (in a re-randomized order) to refresh the materials in the scanner environment.

The critical TNT phase consisted of six runs in a single session, with each run lasting 6.7 minutes and involving the presentation of 16 Think cues and 16 No-Think cues (each cue was presented twice in each run, according to blocked randomization, with the condition assignments for the pairings counterbalanced across participants). Cues from the remaining third of the studied word pairs did not appear during the TNT phase, as they were reserved to provide a baseline measure of memory on the final test, given that they would neither have been suppressed nor retrieved during the TNT phase. As in the practice phase, cue words from the Think condition appeared in green for 3s, indicating that participants were to silently recall the associated target and keep it in mind for the entire time that the cue remained on the screen. For No-Think trials, the cue word appeared in red for an equal duration while participants sought to prevent the associated target word coming into mind; on these No-Think trials, participants were told to directly suppress retrieval to block out the unwanted item, without trying to distract themselves by substituting another word, image, or idea for the unwanted target. Across both conditions, participants were trained to keep their eyes and attention fixed on the presented cues throughout the trial duration. It is worth noting that the word pairs presented in the TNT phase included those that were successfully memorized in the encoding phase, as well as those that were not successfully memorized. Therefore, for the purpose of our analysis, the studied word pairs could be further conditionalized based upon whether participants had successfully recalled the target word in the final test-feedback round of the encoding phase, ensuring that only successfully learned pairs contributed to the analyses we discuss.

#### Memory test phase

After the critical TNT phase, participants completed surprise memory tests for all the studied targets outside the scanner. After asking participants to think back to the original encoding phase (in order to reinstate the encoding phase context), they were asked to recall the targets to 18 filler pairs targets, cued one at a time with the original cue word. This practice test allowed participants to adjust to the instructions to try their best to recall the target words matching the given cues, regardless of what happened in the preceding phase. Participants’ memory for the critical pairings was then tested in a block-randomized fashion with respect to TNT condition, ensuring that the average test position of the Baseline, Think, and No-Think items was equated and that output interference was matched. Each critical target was tested in two ways during this phase, each assessing the accessibility of the learned targets: the same-probe (SP) test, which provided the original cues on the screen for 3.4 s (ISI 0.6 s) as test prompts, then asked subjects to recall the target word. The independent-probe test (IP) test, which measures inhibition in a way that bypasses the original cue-target association and any associative interference the original cue may trigger (Anderson and Green, 2001). Similar to SP test, the IP test provided a category or a semantically related cue on the screen for 3.4 s (ISI 0.6 s) as test prompts, then required subjects to recall the target word fit the cue word. The order of the SP and IP test was counterbalanced across subjects.

### MRI data acquisition

A Siemens 3T scanner (Siemens Magnetom Trio TIM, Erlangen, Germany) was used to collect the functional and structural images from 146 participants. T1-weighted brain anatomical data were collected using a magnetization-prepared rapid gradient echo (MPRAGE) sequence (TR = 1900 ms; FA = 9°; 256 × 256 matrix; TE = 2.52 ms; TI = 900 ms; 176 slices; slice thickness = 1.0 mm; and voxel size = 1 mm × 1 mm × 1 mm). The T2*-weighted functional images were recorded using an echo planar imaging (EPI) sequence (TR = 2000 ms; TE = 30 ms; matrix size = 64 × 64; 32 interleaved 3-mm thick slices; flip angle = 90°; in-plane resolution = 3.4 × 3.4 mm; field of view (FOV) = 220 × 220 mm; and interslice skip = 0.99 mm).

### Image pre-processing

The brain imaging data were preprocessed using the CONN toolbox (version 19.c; Whitfield-Gabrieli and Nieto-Castanon, 2012) in MATLAB vR2018a (The MathWorks, MA, USA). The scans were first coregistered and then resampled to a reference image. Slice-timing correction was used to correct for time shifts by resampling to match the slice time in the middle of each TA. At this stage, we also identified outlier scans, which produced a new reference image by averaging across all scans (except for the outlier scans). The SPM12 unified segmentation and normalization procedure (Ashburner and Friston, 2005) was used to normalize functional images to standard MNI space and segment them into gray matter, white matter, and CSF tissue. We then smoothed the functional data with a Gaussian kernel of 8mm FWHM. However, we did not use the smoothed functional data for our functional connectivity analyses because smoothing may artificially influence individual differences in this prediction analysis (Triana et al., 2020). After preprocessing, the unsmoothed images were denoised using the anatomical component-based correction (aCompCor) method. This procedure removes potential confounding effects, including signals from white matter (WM), cerebrospinal fluid (CSF), and 12 movement parameters (three rotations, three translations, and six first-order temporal derivatives). Importantly, to focus on intrinsic fluctuations in functional connectivity and avoid ambiguous inferences, we also removed the trial-evoked signal (by convolving a box-car model for all task events with canonical hemodynamic response function and plus first and second derivatives; Cole et al., 2019; Goldfarb et al., 2020). We did not apply linear detrending to the time courses because previous research (see Anderson and Hulbert, 2021) indicates that there exist some systematic singal changes over TNT blocks, such as conflict reduction. Temporal band-pass filtering was applied to eliminate low-frequency drift (<0.01Hz). We did not apply global signal (GS) regression because this procedure may cause false negatives correlation in terms of functional connections (Murphy and Fox, 2017), and the GS, itself, may contain neural information (Wen and Liu, 2016).

### Seed regions

The nodes were defined using the default Harvard-Oxford atlas (Tzourio-Mazoyer et al., 2002) applied in CONN toolbox, which including cortical, subcortical and cerebellar areas, parcellates the whole brain into 132 regions. For each participant, the time series of each of the 132 nodes was calculated by averaging the time series of all voxels in that node. We selected both the left and right hippocampus as seed regions and then separately calculated the functional connectivity change between right or left hippocampus with other brain regions (see Generalized psycho-physiological interaction (g-PPI) analysis).

### Generalized psycho-physiological interaction (g-PPI) analysis

The fully preprocessed time course of each node was submitted to g-PPI analysis (using the CONN toolbox; McLaren et al., 2012) to calculate the functional connectivity change between each region (right or left hippocampus) with other brain regions in the No-Think condition compared to the Think condition. The word pairs presented in the TNT phase included those that were successfully memorized in the encoding phase, as well as those that were not successfully memorized. Therefore, we further divided the No-Think and Think conditions into No-Think-Learned, Think-Learned, No-Think-Unlearned and Think-Unlearned conditions. ‘Learned’ represent the word pair was successfully memorized during encoding phase, ‘Unlearned’ represent the word pair that was not successfully memorized during encoding phase. The functional connectivity was computed by using a multiple regression model for the mean time course of right or left hippocampus. The g-PPI model contained three predictors: a) the mean time course for a given seed region; b) the task effects (No-Think-Learned, Think-Learned, No-Think-Unlearned, Think-Unlearned) convolved with a canonical hemodynamic response function; c) the interactions term defined as the product of a) and b). Its objective function can be defined as:

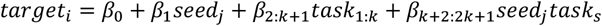

In this formula, target represents the time series of the target node (right or left hippocampus). ‘Seed’ represents the time series of the rest of the brain. The output regression coefficients (β_k+2:2k+1_) of each interaction term (*seed*_*j*_*task*_*s*_) represent the functional connectivity during each condition. The functional connectivity change was computed by subtracting β_Think-Learned_ from β_No-Think-Learned_ and then submitting to the prediction analysis.

### Hippocampal seed connectome-based predictive modeling (seed-CPM)

Functional connectivity change was defined as a subtraction of the two beta matrices: β_No-Think-Learned_ -β_Think-Learned_). Each region’s (right or left hippocampus) connectivity change beta matrix contained 131 features. We separately ran the predictive analysis on the left and right hippocampus. Before the formal prediction analysis, we regressed out the gender, age, and mean framewise displacement from every participant’s functional connectivity change matrix. The leave-one-out cross validation (LOOCV) procedure was used to build the prediction model using the training dataset and test it on the held-out dataset. Briefly, in each fold, N-1 participants served as a training dataset and the left-out participant served as an independent test dataset, a process that was then repeated 134 times. In each fold, we computed the Pearson correlation between each input feature and memory suppression ability, choosing those features that were above-threshold (*p* < 0.05, the observed scores follow the normal distribution, Shapiro-Wilk test: *p* > 0.05). Note that the feature selection step is performed on the training dataset; it is independent of the test data set. Based on the correlation direction, we then divided the chosen features into positive and negative ones and used ridge regression to build the predictive model. The objective function of ridge regression can be defined as:

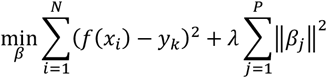

The regularization parameter *λ* is used to shrink the regression coefficient *β*. A large *λ* represents more penalties of variance. Within each fold of the LOOCV, an inner 5-fold cross validation was used to obtain the optimal parameter *λ* from [2^-5^,2^-4^…2^9^,2^10^] (Cui and Gong, 2018). After obtaining predicted behavioral scores for each participant, we computed the model’s performance as the Pearson correlation (r_p_) between participants’ observed and predicted scores. We then used a non-parametric permutation test to test the significance of the correlation coefficient (*r*_p_) by randomly shuffling participants’ observed scores and input beta matrix and then rerunning the whole LOOCV procedure. After 1,000 permutations, we obtained a null distribution of the correlation coefficient between observed and predicted scores. We computed the *p*-value of the true model performance by using the number of null coefficients that were greater than the true r_p_ plus one and then dividing by 1,000. Because a leave-one-out procedure was used in current prediction analysis, the features selected in any given fold varied. To facilitate interpretation of results, in the Results section, we only describe features that appeared in every fold, which we call “consistent features”. The predictive features of right and left hippocampus that selected at least once and their probability been selected are present in Supplementary Table 4-7. The probability represents the percentage of folds in which the feature is identified. The contribution of each consistent feature was calculated as the mean regression coefficient across all folds.

### Model stability

#### Head motion

We took a number of steps to deal with the known potential confound between functional connectivity and head motion (Van Dijk et al., 2012). We removed outlier scans during preprocessing and regressed out mean framewise displacement of each participant’s functional connectivity matrix. We also used partial correlations to measure the predictive power of the model.

#### Cross validation strategy

Although widely used in prediction analysis (Rosenberg et al., 2018; Goldfarb et al., 2020; Beaty et al., 2018), one might be concerned with the LOOCV procedure; because the prediction model built in this way had used almost all the data points, it may effectively reduce biases while also increasing variance of test error (Kohavi, 1995; Varoquaux et al., 2017). From the view of bias-variance trade-off, we also used a 10-fold and 2-fold cross-validation strategy to build the prediction model in a training dataset and test it on the test dataset. 10-fold cross-validation has been shown empirically to not suffer from excessively high variance or very high bias. In other way, 2-fold may produce lower variance, but suffer with high bias. 10-fold and 2-fold are similar to LOOCV procedure, the difference being that, in the 10-fold procedure, we randomly divided the participants into ten groups, nine of which were used as the training set, and the remaining one used as the test set. Similarly, for the 2-fold procedure, half the participants served as the training dataset, while the remaining half served as the test dataset. Because we randomly divided participants into different groups, the prediction performance may differ. Therefore, we repeated the 10-fold or 2-fold prediction test 100 times, and take the mean Pearson correlation between observed scores and predicted scores as the prediction performance and used 1,000 permutations to test for significance.

#### Brain parcellation

Brain parcellation may affect both feature selection and prediction accuracy. Therefore, after obtaining the prediction features defined using the Harvard-Oxford atlas, we then re-ran the procedure instead using the Brainnetome Atlas (Fan et al., 2016). This allowed us to compare whether the predictive features and predictive performance were the same across the two parcellation approaches.

#### Regression method and feature selection threshold

In the main analysis, we used ridge regression to construct a predictive model and with a commonly used feature threshold of *p* < 0.05 (Goldfarb et al., 2020; Rosenberg et al., 2018). To further ensure that our results did not depend on specific regression methods or feature selection thresholds, we also constructed predictive models based on four widely used regression methods: multiple linear regression, Lasso (least absolute shrinkage and selection operator), relevance vector regression, and support vector regression--each under five different feature selection thresholds (0.05 0.01, 0.005, 0.001, 0.0005).

## Results

### Behavioral results

Both the same-probe (SP) and independent-probe (IP) tests were used to assess the final recall percentages for targets from the three TNT conditions (Think, No-Think, and Baseline). Both tests measured a participant’s memory suppression ability by subtracting the recall percentage of the No-Think items from that of the Baseline items. The difference is termed suppression-induced forgetting (SIF). We conditionalized our analysis by considering only those items for which a participant was able to demonstrate successful encoding on their last test-feedback cycle prior to embarking on the TNT phase (Benoit and Anderson, 2012). Because forgetting on the SP test is thought to reflect a mix of inhibition and interference, we chose to use the IP test as a purer measure of inhibition (Anderson and Levy, 2007). Consistent with previous findings and our prior reporting based on this sample of participants (Liu et al., 2021), the conditionalized IP recall of No-Think items (M = 45.47%, SD = 15.73%) was significantly lower than the recall percentage of Baseline (M = 54.75%, SD = 18.25%), yielding a reliable SIF effect (M = 9.25%, SD = 19.98%, one-tailed, *t*_133_ = 5.36, *p* < 0.0001) (see Supplemental Table 1 for a full reporting of means from all tests).

### Prediction of memory suppression ability

We applied seed-CPM to test whether and how memory suppression ability (operationalized as SIF) on the final test was predicted by suppression-related hippocampal functional connectivity changes. We separately ran the predictive analysis on the left and right hippocampus. By computing the Pearson correlation between predicted and observed scores, we found that, compared to Think condition, the functional connectivity change associated with the right hippocampus during the No-Think task significantly predicted memory suppression ability (*r* = 0.36, *p* < 0.05, see Fig. 1a). We then divided the general predictive network into positive and negative networks. Only the negative network predicted participants’ memory suppression ability (positive network: *r* = 0.14, *p* > 0.05; negative network: *r* = 0.35, *p* < 0.05, see Fig. 1b, c). For the left hippocampus, we found that the functional connectivity change did not reliably predict the subject’s memory suppression ability (*r* = 0.21, *p* > 0.05, see Fig. 2a). After we divided the general predictive network into positive and negative networks, the positive networks failed to predict memory suppression ability and the predictive power was marginal in the negative network (positive network: *r* = − 0.05, *p* > 0.05; negative network: *r* = 0.22, *p* = 0.053, see Fig. 2b, c). The p-values presented here were all obtained following 1000 permutation tests. In order to further facilitate the interpretation of the results, we mainly focused on the features that were selected in each and every fold—what we called “consistent” features (Rutherford et al., 2020; Greene et al., 2020; Yang et al., 2021). The predictive features of right and left hippocampus that selected at least once and their probability been selected are present in Supplementary Table 4-7. The consistent right hippocampal negative network revealed that decreased functional connectivity during No-Think trials between the right hippocampus and bilateral intracalcarine cortex, right cuneal cortex, left lingual gyrus, right supracalcarine cortex, right occipital pole, left accumbens and bran-stem predicted higher memory suppression ability (see Fig. 1d). Among these features, left accumbens had the largest weight (see Table 1).

**Fig. 1.**
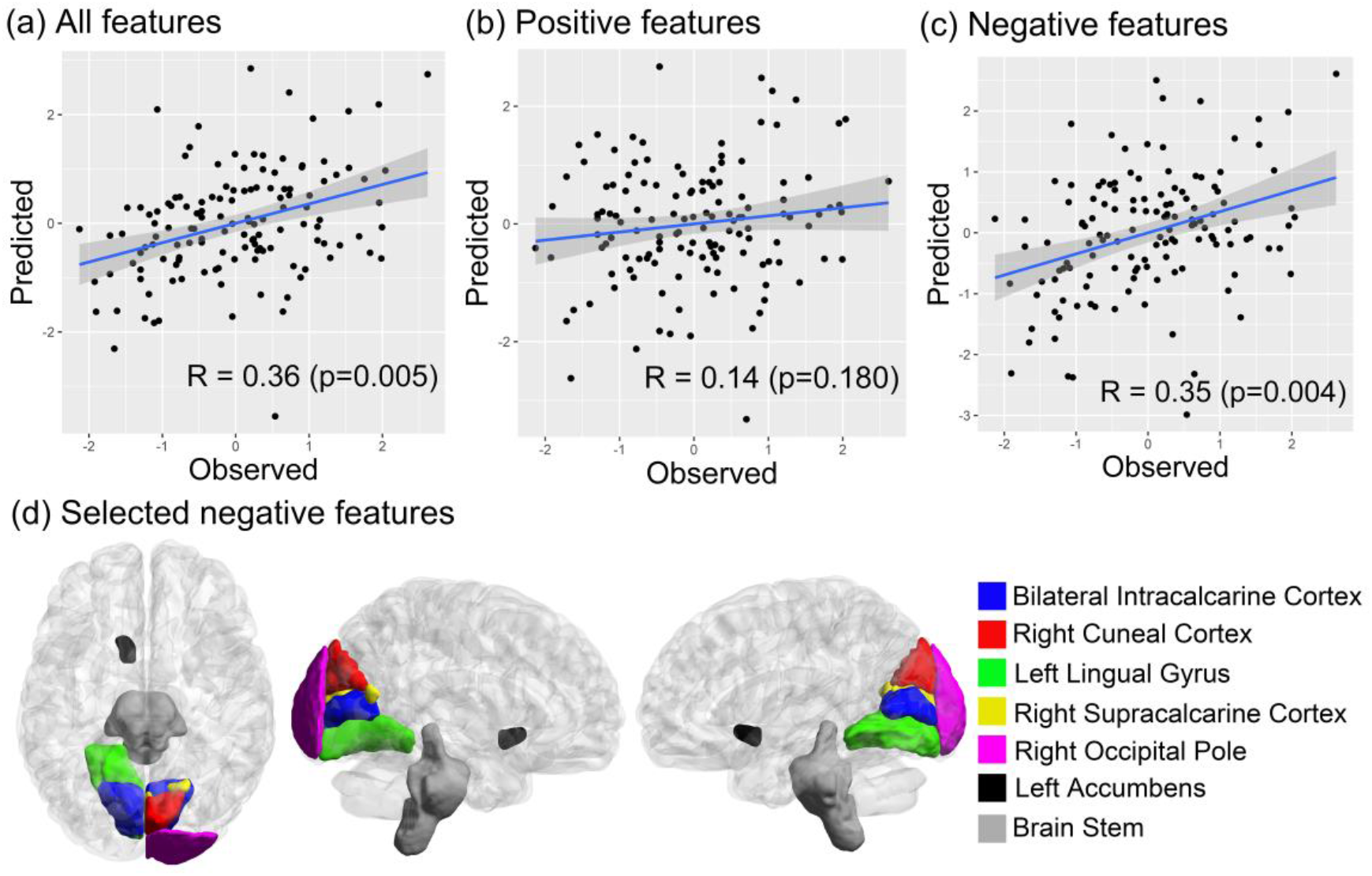
Right Hippocampal functional connectivity change predicts suppression-induced forgetting. The *p*-values here were obtained following 1000 permutation tests. (a) the predictive performance of all selected features; (b) the predictive performance of positive selected features; (c) the predictive performance of negative selected features; (d) The highlighted negative features exist consistently in every cross-validation fold. The brain visualization was generated using BrainNet Viewer (Xia et al., 2013).

**Fig. 2.**
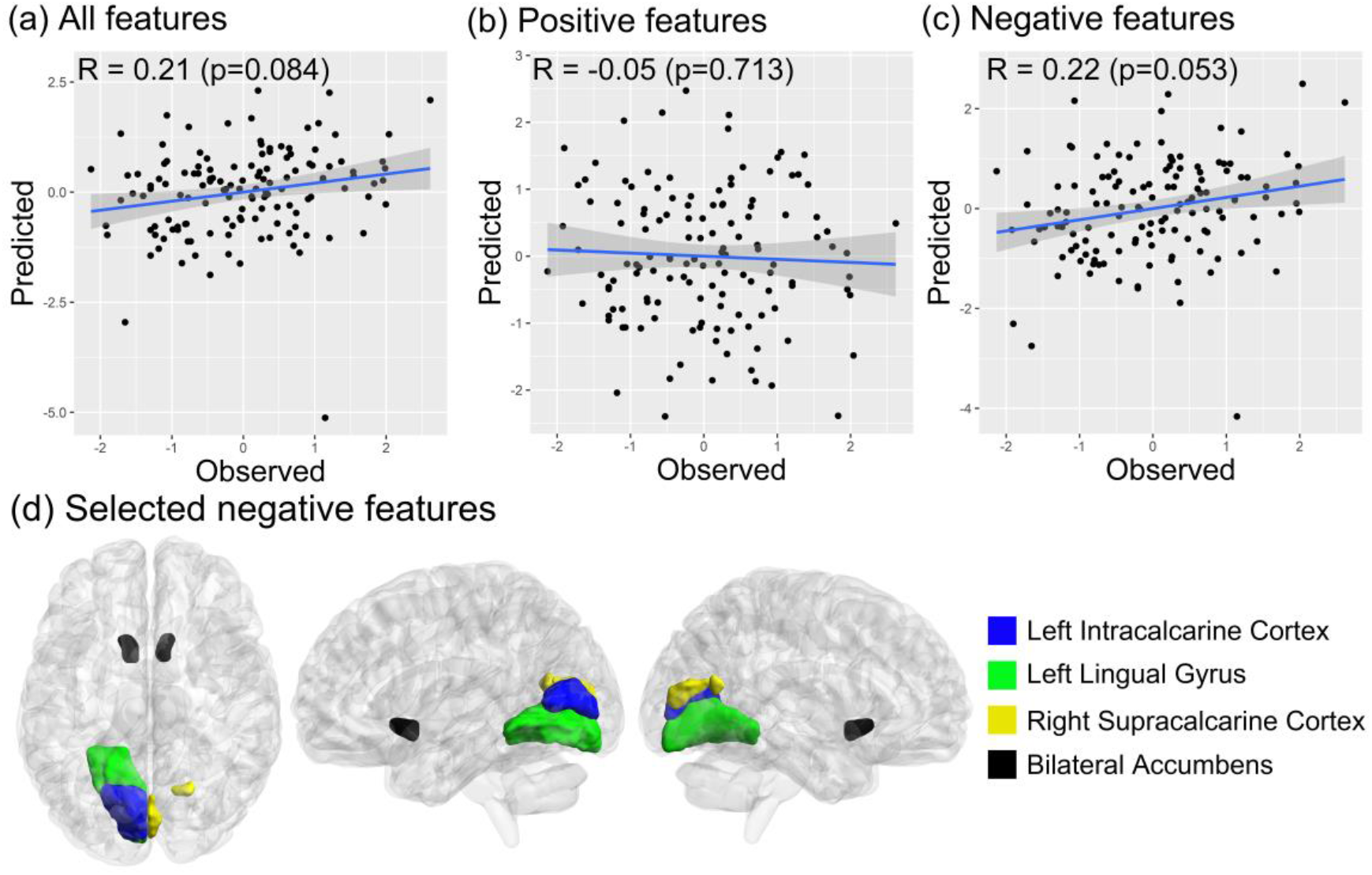
Left Hippocampal functional connectivity change predicts suppression-induced forgetting. The *p*-values here were obtained following 1000 permutation tests. (a) the predictive performance of all selected features; (b) the predictive performance of positive selected features; (c) the predictive performance of negative selected features; (d) The highlighted negative features exist consistently in every cross-validation folds. The brain visualization was generated using BrainNet Viewer (Xia et al., 2013).

**Table 1.** The prediction weight of consistent negative features of right hippocampus.

| Regions | Prediction weight |
| --- | --- |
| Left Intracalcarine Cortex | -0.1305 |
| Right Intracalcarine Cortex | -0.0397 |
| Right Cuneal Cortex | -0.0280 |
| Left Lingual Gyrus | -0.0111 |
| Right Supracalcarine Cortex | -0.0465 |
| Right Occipital Pole | -0.0809 |
| Left Accumbens | -0.1692 |
| Brain Stem | -0.1019 |

The selected negative features had a certain pattern in spatial distribution, in which 75% of the negative features (6/8) were located in the visual cortex, including the Brodmann area 17 and 18 areas. For the left hippocampus, the consistent left hippocampal negative network revealed that decreased functional connectivity during No-Think trials between the left hippocampus and left intracalcarine cortex, left lingual gyrus, right supracalcarine cortex, bilateral accumbens predicted higher memory suppression ability (see Fig. 2d). Among these features, the left accumbens had the largest weight (see Table 2).

**Table 2.**
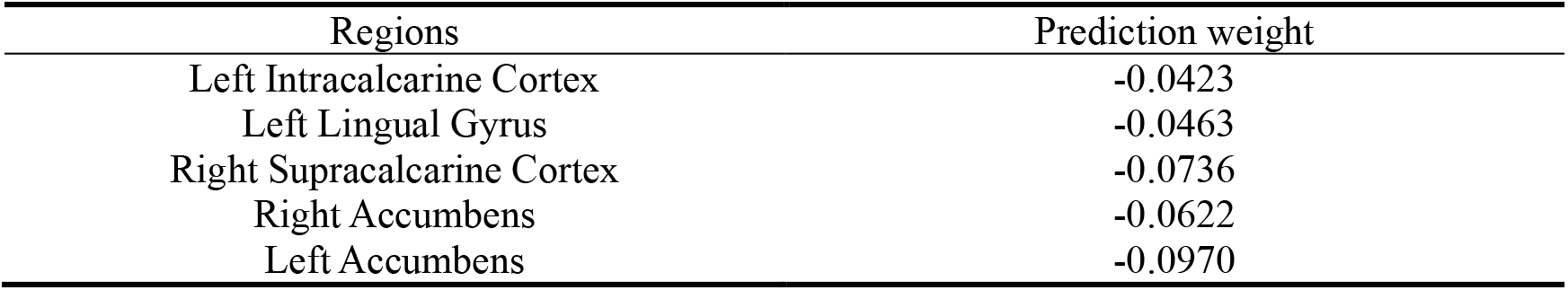
The prediction weight of consistent negative features of left hippocampus.

### The stability of the prediction model

#### Head motion

We applied several methods to confirm that the prediction analysis was not confounded by head motion. First, head motion, calculated as the mean frame-to-frame displacement (mean FD), did not correlate with memory suppression ability (*r* = 0.01, *p* > 0.05). Second, we measured the model performance by comparing the predicted scores to observed scores using partial correlations. After controlling for head motion, the prediction remained unchanged (Right hippocampus: all predictive features: *r* = 0.36, *p* < 0.05, positive predictive features: *r* = 0.14, *p >* 0.05, negative predictive features: *r* = 0.35, *p* < 0.05. Left hippocampus: all predictive features: *r* = 0.21, *p* > 0.05, positive predictive features: *r* = − 0.05, *p >* 0.05, negative predictive features: *r* = 0.22, *p* > 0.05.)

#### Cross-validation strategies

The LOOCV procedure we used in the above analyses may effectively reduce bias while also increasing variance of prediction error. Therefore, we also applied 10-fold and 2-fold cross-validation strategies to verify the stability of the results. For the right hippocampus, the predictive power of all features and negative features remained significant using the 10-fold and 2-fold approaches, despite some expected numeric weakening of the correlation coefficients compared to our primary approach (see Fig. 3). For the 10-fold analysis: *r* = 0.32, *p* < 0.05; positive predictive features: *r* = 0.07, *p* > 0.05; negative predictive features: *r* = 0.32, *p* < 0.05. For 2-fold: all predictive features: *r* = 0.27, *p* < 0.05; positive predictive features: *r* = 0.06, *p* > 0.05; negative predictive features: *r* = 0.28, *p* < 0.05. For the left hippocampus, the functional connectivity change associated with the left hippocampus failed to predict memory suppression ability using the 10-fold and 2-fold approaches. For 10-fold: all predictive features: *r* = 0.10, *p* > 0.05; positive predictive features: *r* = − 0.07, *p* > 0.05; negative predictive features: *r* = 0.16, *p* > 0.05. For 2-fold: all predictive features: *r* = 0.04, *p* > 0.05; positive predictive features: *r* = − 0.03, *p* > 0.05; negative predictive features: *r* = 0.08, *p* > 0.05. Again, the *p*-values presented here were all obtained following 1000 permutation tests.

#### Brain parcellation

In the foregoing analyses, we defined the nodes using the Harvard-Oxford atlas. However, the particulars of the chosen brain parcellation may affect prediction accuracy. Therefore, we applied a different brain parcellation and reran the prediction analyses to determine if the results generalized. The procedure we followed was otherwise the same as above. The results again revealed that the negative features of right hippocampus reliably predicted participants’ memory suppression abilities (*r* = 0.30, *p <* 0.05; see Fig. 4a). Notably, the negative right hippocampal predictive features based on the two parcellation approaches largely overlapped in space, namely the lingual gyrus, cuneus gyrus, and left nucleus accumbens (see Fig. 4b). For the left hippocampus, the negative features failed to predicted subjects’ memory suppression abilities (*r* = 0.10, *p* > 0.05).

**Fig. 3.**
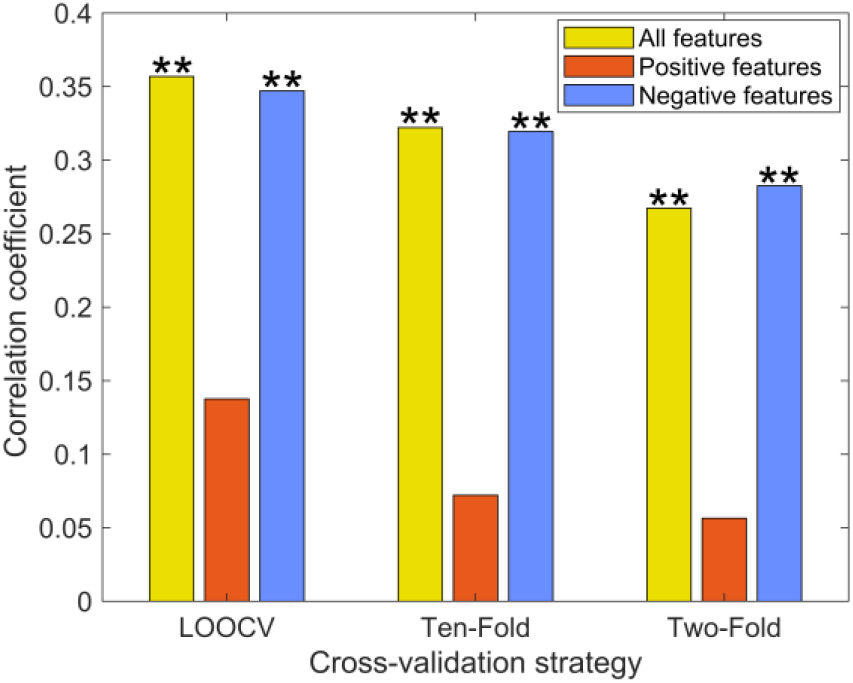
The prediction performance of right hippocampus based on different cross-validation strategies. The *p*-values here were obtained following 1000 permutation tests. * represents *p <* 0.05, ** represents *p <* 0.01; LOOCV: leave-one-out cross validation.

**Fig. 4.**
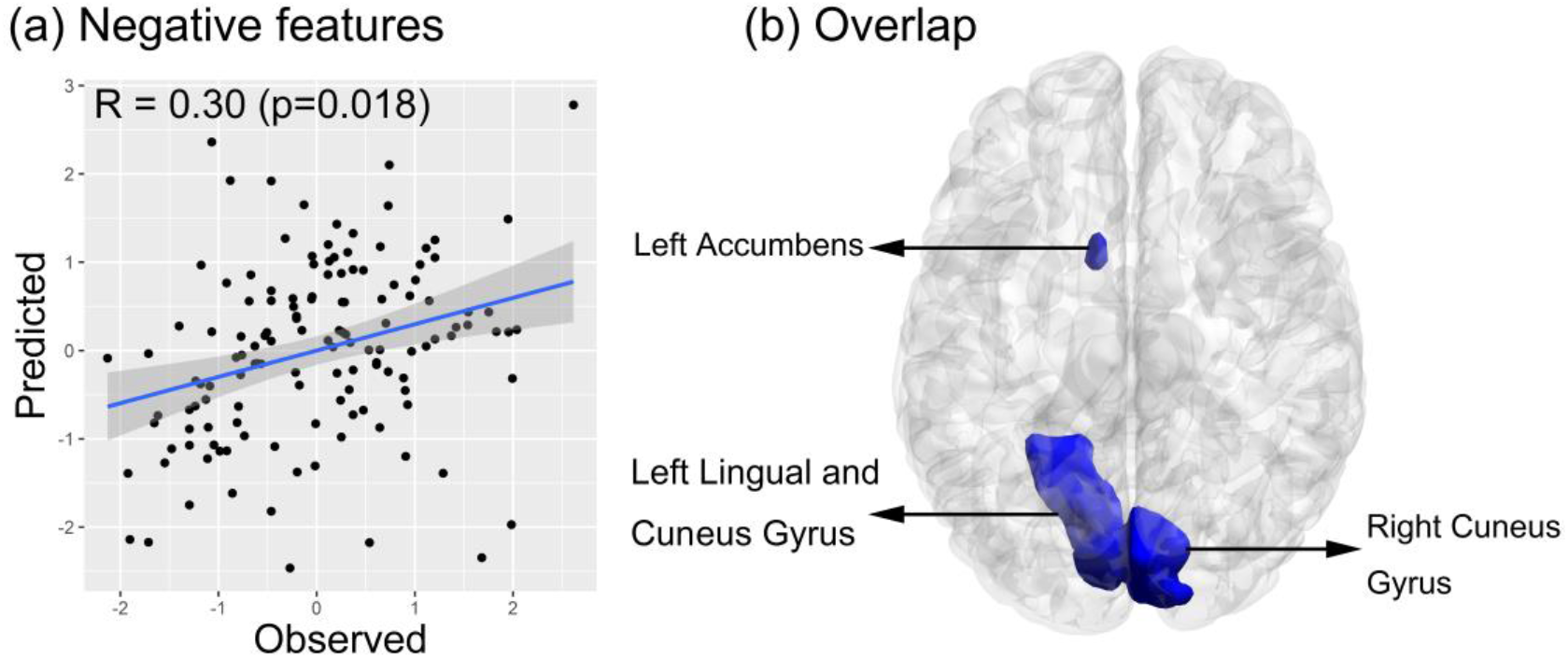
The *p*-values here were obtained following 1000 permutation tests. (a) The prediction performance of right hippocampal negative features based on the Brainnectome atlas; (b) brain map highlighting the right hippocampus consistent negative features that overlapped across the two parcellation approaches (colored in blue); visualization generated with BrainNet Viewer (Xia et al., 2013).

#### Regression methods and feature selection Thresholds

In our primary analyses, ridge regression was used to build the prediction model under the feature selection threshold of 0.05. For validation purposes, we also applied different regression approaches and feature selection thresholds to verify that our right hippocampal results did not depend on any particular regression method or feature selection threshold (see Fig. 5). The results show that different regression methods or feature selection thresholds all produce similar prediction results for right hippocampus. The selected consistent features at different thresholds of right hippocampus are shown in the supplementary Table 2.

**Fig. 5.**
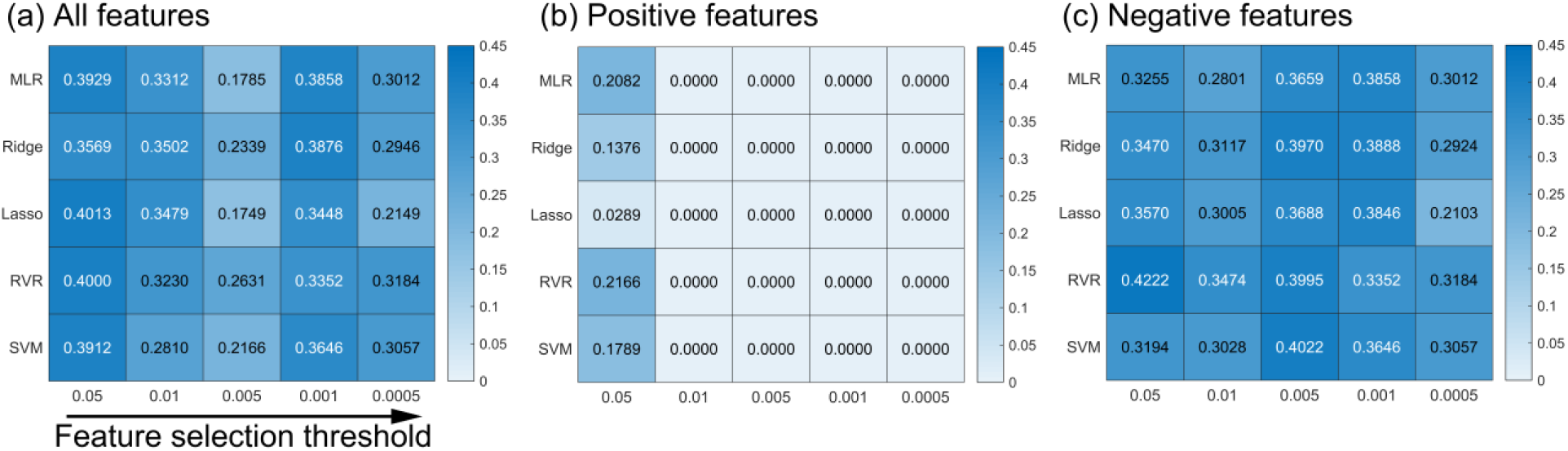
The prediction performance of right hippocampus based on different regression methods. The numbers in the heat map represent the Pearson correlation coefficient between predicted score and observed score. MLR: multiple linear regression; Ridge: ridge regression; Lasso: least absolute shrinkage and selection operator; RVR: relevance vector regression; SVM: support vector regression.

#### Multiple comparisons correction

In the main analysis, we used the left and right hippocampus as nodes and used all the features (both positive and negative) to predict a participant’s memory suppression ability. In other words, 6 (2*3=6) models were trained and tested using LOOCV for each subject. Even we treat the left and right hippocampus as independent region of each other and used the strict Bonferroni correction (0.05/6=0.0083), the primary results of the prediction model built on all or on negative features of the right hippocampus remained significant.

## Discussion

By adopting a g-PPI and seed-CPM analysis approach to neuroimaging data from a large sample of participants engaged in memory suppression, we found that the change in functional connectivity of the right hippocampus during memory suppression predicted participants’ forgetting of suppressed memories on a later recall test. Specifically, decreased functional connectivity of the right hippocampus with visual cortical areas (bilateral intracalcarine cortex, right cuneal cortex, left lingual gyrus, right supracalcarine cortex, right occipital pole), left accumbens and the brainstem predicted superior forgetting ability for suppressed items and formed the negative predictive network. Through several validation tests, we showed that these negative right hippocampal networks were stable across different cross-validation methods, different feature selection thresholds and different regression approaches and did not arise artifactually from head motion. Furthermore, most of the right hippocampal negative features generalized to a different parcellation atlas. Taken together, these results broaden our view of the hippocampus’s role in suppression, moving beyond the simple observation that hippocampal deactivations and negative prefrontal-hippocampal coupling underpin this process (for a recent review, see Anderson and Hulbert, 2021). Our results provide additional support for the notion that, in addition to direct disruption of hippocampally dependent mnemonic processes (for evidence of this, see Hulbert et al., 2016), attempts to stop unwanted memory retrieval depend on reduced communication between the hippocampus and visual cortex.

Many studies have found that memory suppression decreases hippocampal activation (see Anderson and Hulbert, 2021 for a review). Effective connectivity analyses further reveal that reduced activation of these regions is caused in part by top-down control by MFG (Benoit and Anderson, 2012; Benoit et al., 2015; Benoit et al., 2016; Gagnepain et al., 2017; Apšvalka et al., 2022). But most research focuses on a few selected ROIs, such as the right MFG as the source of modulation (Paz-Alonso et al., 2013; Yang et al., 2021) and the hippocampus as the target of down-regulation. However, successful retrieval depends on extensive communication between the hippocampus and many brain regions, including the sensory cortex (Treves and Rolls, 1994; Sutherland and McNaughton, 2000; Schott et al., 2013). Although several studies have documented how suppression affects cortical regions outside the hippocampus that represent the domain-specific content of the to-be-suppressed event (e.g., Gagnepain et al., 2014; Benoit et al., 2015; Gagnepain et al., 2017), no studies have characterized how suppression alters functional connectivity between hippocampus and cortical regions involved in recollection, and the role of such changes play in forgetting. In the present study, we used g-PPI (functional connectivity) and seed-CPM (prediction analysis) to demonstrate that decreased functional connectivity between the right hippocampus and the visual cortex predicts successful forgetting. In addition, we found that other brain regions, such as the nucleus accumbens predict memory suppression ability. Previous research shows that compared to the Think condition, during the No-Think condition, the posterior cortex exhibits a large area of reduced activation, mainly located in the cuneus and the lingual gyrus (Levy and Anderson, 2012; Benoit et al., 2016; Gagnepain et al., 2017; Yang et al., 2020). Our study shows the same pattern: activation is suppressed during the No-Think condition compared to the Think condition, in cuneus and lingual gyrus along with the right hippocampus (see Supplementary Table 3 and Fig. 1). Our results further demonstrated that the visual cortex and hippocampus may not be independent of one another. In addition to the decrease in activation, the functional connectivity between them is also decreased, and this reduction effectively predicted subjects’ later forgetting of the suppressed content.

Of the negative features we identified, the decreased functional connectivity between the right hippocampus and the lingual gyrus and the cuneus gyrus were the most prominent and stable. Both regions send and receive projections to the medial temporal lobes via the inferior longitudinal fasciculus, as demonstrated by diffusion tractography and post-mortem dissection (Palejwala et al., 2021), suggesting a role in memory. Consistent with this possibility, both structural and functional magnetic resonance imaging findings suggest that these components of the ventral visual stream play a role in recollection. For example, greater cortical thickness/grey matter volume in the lingual gyrus predicts better long-term free recall across the lifespan (Walhovd et al., 2006; Kalpouzos et al., 2009). In fMRI studies, lingual gyrus also shows greater univariate activation during episodic retrieval than during encoding (Robinson-Long et al., 2009). Variations in signal in lingual gyrus predict the vividness with which a person can mentally replay a video during episodic retrieval (St-Laurent et al., 2014) and the level of perceptual detail that can be reinstated from previously encoded pictures (McDonough et al., 2014). Wing et al. (2015) found that only the anterior hippocampus and lingual gyrus showed univariate activity at encoding that predicted the degree of item-specific encoding/retrieval match in multivariate analyses, suggesting they helped to establish a recallable and vivid episodic trace. Strikingly, the role of the lingual gyrus may not be restricted to voluntary retrieval but may also extend to involuntary intrusions of unwelcome memories: greater cortical thickness in lingual gyrus predicts participants’ propensity to experience involuntary intrusions during the week after exposure to a traumatic film clip (Gvozdanovic et al., 2020). Collectively, these findings suggest that the visual cortical regions such as the lingual and cuneal gyri contribute to the formation and retrieval of perceptually vivid experiences, as might arise during intrusive memories of trauma. Given these observations, suppressing the retrieval of an intrusive memory might achieve forgetting, in part, by disrupting hippocampal-lingual connectivity that would otherwise support retrieval. Alternatively, retrieval suppression may target both hippocampal and visual cortical activity, and this parallel modulation may be reflected by reduced connectivity (Gagnepain et al., 2014).

We also found that decreased functional connectivity between right hippocampus and left nucleus accumbens predicted superior suppression-induced forgetting of suppressed items. Although retrieval suppression has been found to engage the caudate nucleus and the putamen (Guo et al., 2018), no study has specifically hypothesized a role of the nucleus accumbens in memory suppression. Why this pattern of connectivity predicts suppression-induced forgetting is unclear. Because the anatomical projections linking these structures are unidirectional from the hippocampus to the accumbens (Thierry et al., 2000; Floresco, 2015), it is unlikely that the accumbens could exert an inhibitory impact on hippocampal memory traces to induce forgetting. On the other hand, the nucleus accumbens participates in a network of regions that supports the avoidance of threat, including both active avoidance (taking an action to avoid a threat) and inhibitory avoidance (withholding an action to avoid a threat) (e.g., Levita et al., 2012; Piantadosi et al., 2018). Interestingly, in human imaging studies, inhibitory avoidance (e.g., given a warning cue preceding the appearance of an unpleasant image, withholding a keypress response to prevent the image from appearing) down-regulates BOLD signal in the accumbens, with increasing down-regulation predicting state anxiety (Levita et al., 2012). Because retrieval suppression can be viewed as a form of inhibitory avoidance in which the threat originates from memory, rather than perception, and in which the response is to stop retrieval rather than motor action, the current findings may constitute another example of this intriguing phenomenon. If so, our findings link this form of inhibitory avoidance to later forgetting and to the suppression of hippocampal activity. One simple account of both phenomena is that both retrieval stopping and Levita et al.’s inhibitory avoidance paradigm may lead people to suppress hippocampal activity to prevent recollection of the unwelcome content, disrupting the hippocampus’s connectivity to the nucleus acumbens.

Our demonstration that suppression-induced hippocampal functional connectivity changes could predict participants’ later forgetting is novel in that previous research only achieved this type of prediction using resting-state connectivity of the frontoparietal control networks (Yang et al., 2021). The current findings complement those earlier results. Compared to functional connectivity change induced by a task, resting-state function connectivity tends to be more stable across tasks, and is thus more suitable for predicting individuals’ general competencies (Gratton et al., 2018). However, people face many different tasks in daily life, making it important to dynamically tailor the state of the interconnected brain networks to perform the particular task at hand. Functional connectivity change is more appropriate to predict these dynamic task-related shifts that are missed by resting-state analysis. Previous studies shown that human brain has an intrinsic functional network architecture (Cole et al., 2014) and the functional network is highly similar in different task across subjects (Cole et al., 2014; Krienen et al., 2014; Gratton et al., 2016). Indeed, in present study, we find that the group-averaged hippocampal functional connectivity was highly similar between No-Think and Think condition (right hippocampus: *r* = 0.90, left hippocampus: *r* = 0.90). Gratton et al., (2018) further demonstrated that the intrinsic functional network changes induced by the task are relatively small (∼5%). Our results indicate that, although task-induced changes are small, it also related to partcipants’ cognitive performance. Using functional connectivity change to predict participants’ performance and mental states is a new and promising method. Previous studies have used this method to predict levels of stress from the hippocampal connectivity network and compared this method to others (Goldfarb et al., 2020; Greene et al., 2020). The used of seed connectome-based predictive modeling could also allow researchers to better focus on key brain areas and facilitate the discovery of novel networks involved in a task.

In conclusion, we used machine learning to reveal how the hippocampus communicates functionally with the rest of the brain during memory suppression and in a manner that predicts behavior on a later memory test. Overall, our results suggest that active suppression yields forgetting through decreases in the functional connectivity between the right hippocampus with visual cortex (lingual gyrus and cuneus gyrus) and nucleus acumbens. The identified hippocampal networks provide insight into neurobiological mechanisms supporting active forgetting, which could, in turn, have profound implications for mental disorders characterized by intrusive memories.

## References

Albers, A.M., Kok, P., Toni, I., Dijkerman, H.C., De Lange, F., 2013. Shared representations for working memory and mental imagery in early visual cortex. Current Biology 23, 1427–1431. https://doi.org/10.1016/j.cub.2013.05.065

Anderson, M.C., Green, C., 2001. Suppressing unwanted memories by executive control. Nature 410, 366–369. https://doi.org/10.1038/35066572

Anderson, M.C., Levy, B.J., 2007. Theoretical issues in inhibition: Insights from research on human memory. https://doi.org/10.1037/11587-005

Anderson, M.C., Huddleston, E., 2012. Towards a cognitive and neurobiological model of motivated forgetting. True false recovered memories, 53–120. https://doi.org/10.1007/978-1-4614-1195-6_3

Anderson, M.C., Hanslmayr, S., 2014. Neural mechanisms of motivated forgetting. Trends in cognitive sciences 18, 279–292. https://doi.org/10.1016/j.tics.2014.03.002

Anderson, M.C., Hulbert, J.C., 2021. Active forgetting: Adaptation of memory by prefrontal control. Annual Review of Psychology 72, 1–36. https://doi.org/10.1146/annurev-psych-072720-094140

Anderson, M.C., Ochsner, K.N., Kuhl, B., Cooper, J., Robertson, E., Gabrieli, S.W., Glover, G.H., Gabrieli, J.D., 2004. Neural systems underlying the suppression of unwanted memories. Science 303, 232–235. DOI: 10.1126/science.1089504

Apšvalka, D., Ferreira, C.S., Schmitz, T.W., Rowe, J.B., Anderson, M., 2022. Dynamic targeting enables domain-general inhibitory control over action and thought by the prefrontal cortex. Nature Communications 13, 1–21. https://doi.org/10.1038/s41467-021-27926-w

Ashburner, J., Friston, K.J., 2005. Unified segmentation. Neuroimage 26, 839–851. https://doi.org/10.1016/j.neuroimage.2005.02.018

Backus, A.R., Bosch, S.E., Ekman, M., Grabovetsky, A.V., Doeller, C.F., 2016. Mnemonic convergence in the human hippocampus. Nature Communications 7, 1–9. https://doi.org/10.1038/ncomms11991

Beaty, R.E., Kenett, Y.N., Christensen, A.P., Rosenberg, M.D., Benedek, M., Chen, Q., Fink, A., Qiu, J., Kwapil, T.R., Kane, M.J., 2018. Robust prediction of individual creative ability from brain functional connectivity. Proceedings of the National Academy of Sciences 115, 1087–1092. https://doi.org/10.1073/pnas.1713532115

Benoit, R.G., Anderson, M.C., 2012. Opposing mechanisms support the voluntary forgetting of unwanted memories. Neuron 76, 450–460. https://doi.org/10.1016/j.neuron.2012.07.025

Benoit, R.G., Davies, D.J., Anderson, M.C., 2016. Reducing future fears by suppressing the brain mechanisms underlying episodic simulation. Proceedings of the National Academy of Sciences 113, E8492–E8501. https://doi.org/10.1073/pnas.1606604114

Benoit, R.G., Hulbert, J.C., Huddleston, E., Anderson, M.C., 2015. Adaptive top–down suppression of hippocampal activity and the purging of intrusive memories from consciousness. Journal of cognitive neuroscience 27, 96–111. https://doi.org/10.1162/jocn_a_00696

Cole, M.W., Ito, T., Schultz, D., Mill, R., Chen, R., Cocuzza, C., 2019. Task activations produce spurious but systematic inflation of task functional connectivity estimates. Neuroimage 189, 1–18. https://doi.org/10.1016/j.neuroimage.2018.12.054

Cole, M.W., Bassett, D.S., Power, J.D., Braver, T.S., Petersen, S.E., 2014. Intrinsic and task-evoked network architectures of the human brain. Neuron 83, 238–251. https://doi.org/10.1016/j.neuron.2014.05.014

Cui, Z., Gong, G., 2018. The effect of machine learning regression algorithms and sample size on individualized behavioral prediction with functional connectivity features. Neuroimage 178, 622–637. https://doi.org/10.1016/j.neuroimage.2018.06.001

Depue, B.E., 2012. A neuroanatomical model of prefrontal inhibitory modulation of memory retrieval. Neuroscience & Biobehavioral Reviews 36, 1382–1399. https://doi.org/10.1016/j.neubiorev.2012.02.012

Depue, B.E., Curran, T., Banich, M.T., 2007. Prefrontal regions orchestrate suppression of emotional memories via a two-phase process. Science 317, 215–219. DOI: 10.1126/science.1139560

Fan, L., Li, H., Zhuo, J., Zhang, Y., Wang, J., Chen, L., Yang, Z., Chu, C., Xie, S., Laird, A.R., 2016. The human brainnetome atlas: a new brain atlas based on connectional architecture. Cerebral Cortex 26, 3508–3526. https://doi.org/10.1093/cercor/bhw157

Fawcett, J.M., Hulbert, J.C., 2020. The many faces of forgetting: Toward a constructive view of forgetting in everyday life. Journal of Applied Research in Memory Cognition 9, 1–18. https://doi.org/10.1016/j.jarmac.2019.11.002

Floresco, S.B., 2015. The nucleus accumbens: an interface between cognition, emotion, and action. Annual Review of Psychology 66, 25–52. https://doi.org/10.1146/annurev-psych-010213-115159

Gagnepain, P., Henson, R.N., Anderson, M.C., 2014. Suppressing unwanted memories reduces their unconscious influence via targeted cortical inhibition. Proceedings of the National Academy of Sciences 111, E1310–E1319. https://doi.org/10.1073/pnas.1311468111

Gagnepain, P., Hulbert, J., Anderson, M.C., 2017. Parallel regulation of memory and emotion supports the suppression of intrusive memories. Journal of Neuroscience 37, 6423–6441. https://doi.org/10.1523/JNEUROSCI.2732-16.2017

Geib, B.R., Stanley, M.L., Wing, E.A., Laurienti, P.J., Cabeza, R., 2017. Hippocampal contributions to the large-scale episodic memory network predict vivid visual memories. Cerebral Cortex 27, 680–693. https://doi.org/10.1093/cercor/bhv272

Goldfarb, E.V., Rosenberg, M.D., Seo, D., Constable, R.T., Sinha, R., 2020. Hippocampal seed connectome-based modeling predicts the feeling of stress. Nature Communications 11, 1–10. https://doi.org/10.1038/s41467-020-16492-2

Gratton, C., Laumann, T.O., Nielsen, A.N., Greene, D.J., Gordon, E.M., Gilmore, A.W., Nelson, S.M., Coalson, R.S., Snyder, A.Z., Schlaggar, B.L., 2018. Functional brain networks are dominated by stable group and individual factors, not cognitive or daily variation. Neuron 98, 439–452. e435. https://doi.org/10.1016/j.neuron.2018.03.035

Gratton, C., Laumann, T.O., Gordon, E.M., Adeyemo, B., Petersen, S.E., 2016. Evidence for two independent factors that modify brain networks to meet task goals. Cell reports 17, 1276–1288. https://doi.org/10.1016/j.celrep.2016.10.002

Greene, A.S., Gao, S., Noble, S., Scheinost, D., Constable, R.T., 2020. How tasks change whole-brain functional organization to reveal brain-phenotype relationships. Cell reports 32, 108066. https://doi.org/10.1016/j.celrep.2020.108066

Guo, Y., Schmitz, T.W., Mur, M., Ferreira, C.S., Anderson, M.C., 2018. A supramodal role of the basal ganglia in memory and motor inhibition: Meta-analytic evidence. Neuropsychologia 108, 117–134. https://doi.org/10.1016/j.neuropsychologia.2017.11.033

Guzman, S.J., Schlögl, A., Frotscher, M., Jonas, P., 2016. Synaptic mechanisms of pattern completion in the hippocampal CA3 network. Science 353, 1117–1123. DOI: 10.1126/science.aaf1836

Gvozdanovic, G., Stämpfli, P., Seifritz, E., Rasch, B., 2020. Structural brain differences predict early traumatic memory processing. Psychophysiology 57, e13354. https://doi.org/10.1111/psyp.13354

Hindy, N.C., Ng, F.Y., Turk-Browne, N.B., 2016. Linking pattern completion in the hippocampus to predictive coding in visual cortex. Nature neuroscience 19, 665–667. https://doi.org/10.1038/nn.4284

Horner, A.J., Bisby, J.A., Bush, D., Lin, W.-J., Burgess, N., 2015. Evidence for holistic episodic recollection via hippocampal pattern completion. Nature Communications 6, 1–11. https://doi.org/10.1038/ncomms8462

Huang, C.-C., Rolls, E.T., Hsu, C.-C.H., Feng, J., Lin, C.-P., 2021. Extensive cortical connectivity of the human hippocampal memory system: beyond the “what” and “where” dual stream model. Cerebral Cortex 31, 4652–4669. https://doi.org/10.1093/cercor/bhab113

Hulbert, J.C., Henson, R.N., Anderson, M.C., 2016. Inducing amnesia through systemic suppression. Nature Communications 7, 1–9. https://doi.org/10.1038/ncomms11003

Kalpouzos, G., Chételat, G., Baron, J.-C., Landeau, B., Mevel, K., Godeau, C., Barré, L., Constans, J.-M., Viader, F., Eustache, F., 2009. Voxel-based mapping of brain gray matter volume and glucose metabolism profiles in normal aging. Neurobiology of aging 30, 112–124. https://doi.org/10.1016/j.neurobiolaging.2007.05.019

Kohavi, R., 1995. A study of cross-validation and bootstrap for accuracy estimation and model selection. Ijcai. Montreal, Canada, pp. 1137–1145.

Krienen, F.M., Yeo, B.T., Buckner, R.L., 2014. Reconfigurable task-dependent functional coupling modes cluster around a core functional architecture. Philosophical Transactions of the Royal Society B: Biological Sciences 369, 20130526. https://doi.org/10.1098/rstb.2013.0526

Levita, L., Hoskin, R., Champi, S., 2012. Avoidance of harm and anxiety: a role for the nucleus accumbens. Neuroimage 62, 189–198. https://doi.org/10.1016/j.neuroimage.2012.04.059

Levy, B.J., Anderson, M.C., 2012. Purging of memories from conscious awareness tracked in the human brain. Journal of Neuroscience 32, 16785–16794. https://doi.org/10.1523/JNEUROSCI.2640-12.2012

Liu, P., Hulbert, J.C., Yang, W., Guo, Y., Qiu, J., Anderson, M.C., 2021. Task compliance predicts suppression-induced forgetting in a large sample. Scientific reports 11, 1–13. https://doi.org/10.1038/s41598-021-99806-8

Marsh, L., Anderson, M., 2020. Inhibition as a cause of forgetting. Oxford University Press Oxford.

Mary, A., Dayan, J., Leone, G., Postel, C., Fraisse, F., Malle, C., Vallée, T., Klein-Peschanski, C., Viader, F., De la Sayette, V., 2020. Resilience after trauma: The role of memory suppression. Science 367. DOI: 10.1126/science.aay8477

McDonough, I.M., Cervantes, S.N., Gray, S.J., Gallo, D.A., 2014. Memory’s aging echo: Age-related decline in neural reactivation of perceptual details during recollection. Neuroimage 98, 346–358. https://doi.org/10.1016/j.neuroimage.2014.05.012

McLaren, D.G., Ries, M.L., Xu, G., Johnson, S.C., 2012. A generalized form of context-dependent psychophysiological interactions (gPPI): a comparison to standard approaches. Neuroimage 61, 1277–1286. https://doi.org/10.1016/j.neuroimage.2012.03.068

Murphy, K., Fox, M.D., 2017. Towards a consensus regarding global signal regression for resting state functional connectivity MRI. Neuroimage 154, 169–173. https://doi.org/10.1016/j.neuroimage.2016.11.052

Nørby, S., 2015. Why forget? On the adaptive value of memory loss. Perspectives on Psychological Science 10, 551–578. https://doi.org/10.1177/1745691615596787

Nyberg, L., Habib, R., McIntosh, A.R., Tulving, E., 2000. Reactivation of encoding-related brain activity during memory retrieval. Proceedings of the National Academy of Sciences 97, 11120–11124. https://doi.org/10.1073/pnas.97.20.11120

Paz-Alonso, P.M., Bunge, S.A., Anderson, M.C., Ghetti, S., 2013. Strength of coupling within a mnemonic control network differentiates those who can and cannot suppress memory retrieval. Journal of Neuroscience 33, 5017–5026. https://doi.org/10.1523/JNEUROSCI.3459-12.2013

Palejwala, A.H., Dadario, N.B., Young, I.M., O’Connor, K., Briggs, R.G., Conner, A.K., O’Donoghue, D.L., Sughrue, M.E., 2021. Anatomy and white matter connections of the lingual gyrus and cuneus. World Neurosurgery 151, e426–e437. https://doi.org/10.1016/j.wneu.2021.04.050

Piantadosi, P.T., Yeates, D.C., Floresco, S.B., 2018. Cooperative and dissociable involvement of the nucleus accumbens core and shell in the promotion and inhibition of actions during active and inhibitory avoidance. Neuropharmacology 138, 57–71. https://doi.org/10.1016/j.neuropharm.2018.05.028

Ranganath, C., Heller, A., Cohen, M.X., Brozinsky, C.J., Rissman, J., 2005. Functional connectivity with the hippocampus during successful memory formation. Hippocampus 15, 997–1005. https://doi.org/10.1002/hipo.20141

Robinson-Long, M., Eslinger, P.J., Wang, J., Meadowcroft, M., Yang, Q., 2009. Functional MRI evidence for distinctive binding and consolidation pathways for face-name associations: analysis of activation maps and BOLD response amplitudes. Topics in Magnetic Resonance Imaging 20, 271–278. DOI: 10.1097/RMR.0b013e3181e8f1f9

Rolls, E., 2013. The mechanisms for pattern completion and pattern separation in the hippocampus. Frontiers in systems neuroscience 7, 74. https://doi.org/10.3389/fnsys.2013.00074

Rosenberg, M.D., Hsu, W.-T., Scheinost, D., Todd Constable, R., Chun, M.M., 2018. Connectome-based models predict separable components of attention in novel individuals. Journal of cognitive neuroscience 30, 160–173. https://doi.org/10.1162/jocn_a_01197

Rosenthal, C.R., Andrews, S.K., Antoniades, C.A., Kennard, C., Soto, D., 2016. Learning and recognition of a nonconscious sequence of events in human primary visual cortex. Current Biology 26, 834–841. https://doi.org/10.1016/j.cub.2016.01.040

Rutherford, H.J., Potenza, M.N., Mayes, L.C., Scheinost, D., 2020. The application of connectome-based predictive modeling to the maternal brain: implications for mother–infant bonding. Cerebral Cortex 30, 1538–1547. https://doi.org/10.1093/cercor/bhz185

Schott, B.H., Wüstenberg, T., Wimber, M., Fenker, D.B., Zierhut, K.C., Seidenbecher, C.I., Heinze, H.J., Walter, H., Düzel, E., Richardson-Klavehn, A., 2013. The relationship between level of processing and hippocampal–cortical functional connectivity during episodic memory formation in humans. Human brain mapping 34, 407–424. https://doi.org/10.1002/hbm.21435

Shen, X., Finn, E.S., Scheinost, D., Rosenberg, M.D., Chun, M.M., Papademetris, X., Constable, R.T., 2017. Using connectome-based predictive modeling to predict individual behavior from brain connectivity. nature protocols 12, 506–518. https://doi.org/10.1038/nprot.2016.178

Spiers, H.J., Maguire, E.A., Burgess, N., 2001. Hippocampal amnesia. Neurocase 7, 357–382. https://doi.org/10.1076/neur.7.5.357.16245

St-Laurent, M., Abdi, H., Bondad, A., Buchsbaum, B.R., 2014. Memory reactivation in healthy aging: evidence of stimulus-specific dedifferentiation. Journal of Neuroscience 34, 4175–4186. https://doi.org/10.1523/JNEUROSCI.3054-13.2014

Stramaccia, D.F., Meyer, A.-K., Rischer, K.M., Fawcett, J.M., Benoit, R.G., 2021. Memory suppression and its deficiency in psychological disorders: A focused meta-analysis. Journal of Experimental Psychology: General 150, 828. https://doi.org/10.1037/xge0000971

Sutherland, G.R., McNaughton, B., 2000. Memory trace reactivation in hippocampal and neocortical neuronal ensembles. Current opinion in neurobiology 10, 180–186. https://doi.org/10.1016/S0959-4388(00)00079-9

Thierry, A.M., Gioanni, Y., Dégénétais, E., Glowinski, J., 2000. Hippocampo-prefrontal cortex pathway: Anatomical and electrophysiological characteristics. Hippocampus 10, 411–419. https://doi.org/10.1002/10981063(2000)10:4<411::AID-HIPO7>3.0.CO;2-A

Treves, A., Rolls, E.T., 1994. Computational analysis of the role of the hippocampus in memory. Hippocampus 4, 374–391. https://doi.org/10.1002/hipo.450040319

Triana, A.M., Glerean, E., Saramäki, J., Korhonen, O., 2020. Effects of spatial smoothing on group-level differences in functional brain networks. Network Neuroscience 4, 556–574. https://doi.org/10.1162/netn_a_00132

Tzourio-Mazoyer, N., Landeau, B., Papathanassiou, D., Crivello, F., Etard, O., Delcroix, N., Mazoyer, B., Joliot, M., 2002. Automated anatomical labeling of activations in SPM using a macroscopic anatomical parcellation of the MNI MRI single-subject brain. Neuroimage 15, 273–289. https://doi.org/10.1006/nimg.2001.0978

Van Dijk, K.R., Sabuncu, M.R., Buckner, R.L., 2012. The influence of head motion on intrinsic functional connectivity MRI. Neuroimage 59, 431–438. https://doi.org/10.1016/j.neuroimage.2011.07.044

Varoquaux, G., Raamana, P.R., Engemann, D.A., Hoyos-Idrobo, A., Schwartz, Y., Thirion, B., 2017. Assessing and tuning brain decoders: cross-validation, caveats, and guidelines. Neuroimage 145, 166–179. https://doi.org/10.1016/j.neuroimage.2016.10.038

Waldhauser, G.T., Braun, V., Hanslmayr, S., 2016. Episodic memory retrieval functionally relies on very rapid reactivation of sensory information. Journal of Neuroscience 36, 251–260. https://doi.org/10.1523/JNEUROSCI.2101-15.2016

Walhovd, K.B., Fjell, A.M., Dale, A.M., Fischl, B., Quinn, B.T., Makris, N., Salat, D., Reinvang, I., 2006. Regional cortical thickness matters in recall after months more than minutes. Neuroimage 31, 1343–1351. https://doi.org/10.1016/j.neuroimage.2006.01.011

Wen, H., Liu, Z., 2016. Broadband electrophysiological dynamics contribute to global resting-state fMRI signal. Journal of Neuroscience 36, 6030–6040. https://doi.org/10.1523/JNEUROSCI.0187-16.2016

Wheeler, M.E., Petersen, S.E., Buckner, R.L., 2000. Memory’s echo: vivid remembering reactivates sensory-specific cortex. Proceedings of the National Academy of Sciences 97, 11125–11129. https://doi.org/10.1073/pnas.97.20.11125

Whitfield-Gabrieli, S., Nieto-Castanon, A., 2012. Conn: a functional connectivity toolbox for correlated and anticorrelated brain networks. Brain connectivity 2, 125–141. https://doi.org/10.1089/brain.2012.0073

Wing, E.A., Ritchey, M., Cabeza, R., 2015. Reinstatement of individual past events revealed by the similarity of distributed activation patterns during encoding and retrieval. Journal of cognitive neuroscience 27, 679–691. https://doi.org/10.1162/jocn_a_00740

Wolosin, S.M., Zeithamova, D., Preston, A.R., 2012. Reward modulation of hippocampal subfield activation during successful associative encoding and retrieval. Journal of cognitive neuroscience 24, 1532–1547. https://doi.org/10.1162/jocn_a_00237

Xia, M., Wang, J., He, Y., 2013. BrainNet Viewer: a network visualization tool for human brain connectomics. PloS one 8, e68910. https://doi.org/10.1371/journal.pone.0068910

Yang, W., Liu, P., Zhuang, K., Wei, D., Anderson, M.C., Qiu, J., 2020. Behavioral and neural correlates of memory suppression in subthreshold depression. Psychiatry Research: Neuroimaging 297, 111030. https://doi.org/10.1016/j.pscychresns.2020.111030

Yang, W., Zhuang, K., Liu, P., Guo, Y., Chen, Q., Wei, D., Qiu, J., 2021. Memory Suppression Ability can be Robustly Predicted by the Internetwork Communication of Frontoparietal Control Network. Cerebral Cortex 31, 3451–3461. https://doi.org/10.1093/cercor/bhab024

Zhu, Z., Wang, Y., 2021. Forgetting unrelated episodic memories through suppression-induced amnesia. Journal of Experimental Psychology: General 150, 401. https://doi.org/10.1037/xge0000782.

