## Supplementary for "Reduced Hippocampal-Cortical Connectivity During Memory Suppression Predicts the Ability to Forget Unwanted Memories"

**Supplementary materials**

**Tables**

**Table 1.** The recall percentages observed on the Same-Probe (SP) and the Independent-Probe (IP) tests. Conditionalized data refers to final recall scores, restricted to items that were demonstrably learned during study; unconditionalized data includes all items. Values in brackets represent the 95% confidence intervals for the marginal means.

| **Condition** | **Think** | **No-Think** | **Baseline** |
| --- | --- | --- | --- |
| **Same-Probe test** | | | |
| Conditionalized | 92% [90 94] | 83% [80 86] | 89% [87 91] |
| Unconditionalized | 81% [79 84] | 72% [68 75] | 76% [73 79] |
| **Independent-Probe test** | | | |
| Conditionalized | 43% [40 46] | 46% [43 48] | 55% [52 58] |
| Unconditionalized | 39% [36 41] | 40% [37 42] | 47% [44 50] |

**Table 2.** The consistent negative features (selected in every fold using leave-one-out cross-validation) of the right hippocampus under different features selection threshold (0.01 0.005 0.001 0.0005). The stars represent the features selected under this threshold.

| Regions | 0.01 | 0.005 | 0.001 | 0.0005 |
| --- | --- | --- | --- | --- |
| Left Intracalcarine Cortex | * | * | * | * |
| Right Intracalcarine Cortex | * | * |  |  |
| Right Cuneal Cortex |  |  |  |  |
| Left Lingual Gyrus | * |  |  |  |
| Right Supracalcarine Cortex | * | * |  |  |
| Right Occipital Pole |  |  |  |  |
| Left Accumbens | * | * |  |  |
| Brain Stem |  |  |  |  |

**Table 3.** The averaged activation across all voxels within the visual cortex (selected as negative consistent features of right hippocampus) and the averaged activation across all voxels within the right hippocampus. Negative T values represent lower activation in No-Think trials compared to Think trials.

| Regions | T values (two-tailed) | P values |
| --- | --- | --- |
| Right Hippocampus  Visual Cortex | -5.5  -12.1 | 1.8e-07  3.0e-23 |

**Table 4.** The negative features of the right hippocampus that were selected at least once and the percentage of folds in which they were selected.

| Regions | Percentage |
| --- | --- |
| Left Intracalcarine Cortex | 100.0％ |
| Right Intracalcarine Cortex | 100.0％ |
| Right Cuneal Cortex | 100.0％ |
| Left Cuneal Cortex | 6.0％ |
| Right Parahippocampal Gyrus (posterior division) | 0.8％ |
| Right Lingual Gyrus | 99.3％ |
| Left Lingual Gyrus | 100.0％ |
| Right Occipital Fusiform Gyrus | 5.2％ |
| Right Supracalcarine Cortex | 100.0％ |
| Right Occipital Pole | 100.0％ |
| Right Thalamus | 8.2％ |
| Left Thalamus | 0.8％ |
| Right Accumbens | 99.3％ |
| Left Accumbens | 100.0％ |
| Brain-Stem | 100.0％ |
| Left Cereb10 | 2.2％ |
| Vermis 7 | 88.1％ |

**Table 5.** The positive features of the right hippocampus that were selected at least once and the percentage of folds in which they were selected.

| Regions | Percentage |
| --- | --- |
| Left Frontal Pole | 17.2％ |
| Left Middle Temporal Gyrus (posterior division) | 0.8％ |
| Left Middle Temporal Gyrus (temporooccipital part) | 1.5％ |
| Right Inferior Temporal Gyrus (posterior division) | 6.7％ |
| Left Inferior Temporal Gyrus (posterior division) | 98.5％ |
| Right Postcentral Gyrus | 0.8％ |
| Right Superior Parietal Lobule | 100.0％ |
| Right Angular Gyrus | 0.8％ |
| Left Angular Gyrus | 99.3％ |
| Right Central Opercular Cortex | 1.5％ |
| Right Parietal Operculum Cortex | 3.0％ |
| Right Heschl's Gyrus | 33.6％ |
| Left Heschl's Gyrus | 100.0％ |

**Table 6.** The negative features of left hippocampus that were selected at least once and the percentage of folds in which they were selected.

| Regions | Percentage |
| --- | --- |
| Right Intracalcarine Cortex | 83.6％ |
| Left Intracalcarine Cortex | 100.0％ |
| Right Cuneal Cortex | 15.7％ |
| Left Lingual Gyrus | 100.0％ |
| Left Occipital Fusiform Gyrus | 0.8％ |
| Right Supracalcarine Cortex | 100.0％ |
| Right Accumbens | 100.0％ |
| Left Accumbens | 100.0％ |
| Brain-Stem | 73.1％ |
| Left Cereb10 | 99.3％ |
| Vermis 6 | 0.8％ |

**Table 7.** The positive features of left hippocampus that were selected at least once and the percentage of folds in which they were selected.

| Regions | Percentage |
| --- | --- |
| Right Frontal Pole | 5.2％ |
| Left Frontal Pole | 3.0％ |
| Right Middle Temporal Gyrus (anterior division) | 0.8％ |
| Right Middle Temporal Gyrus (temporooccipital part) | 1.5％ |
| Right Inferior Temporal Gyrus (posterior division) | 3.0％ |
| Right Superior Parietal Lobule | 100.0％ |
| Right Lateral Occipital Cortex (superior division) | 6.0％ |

**Figures**

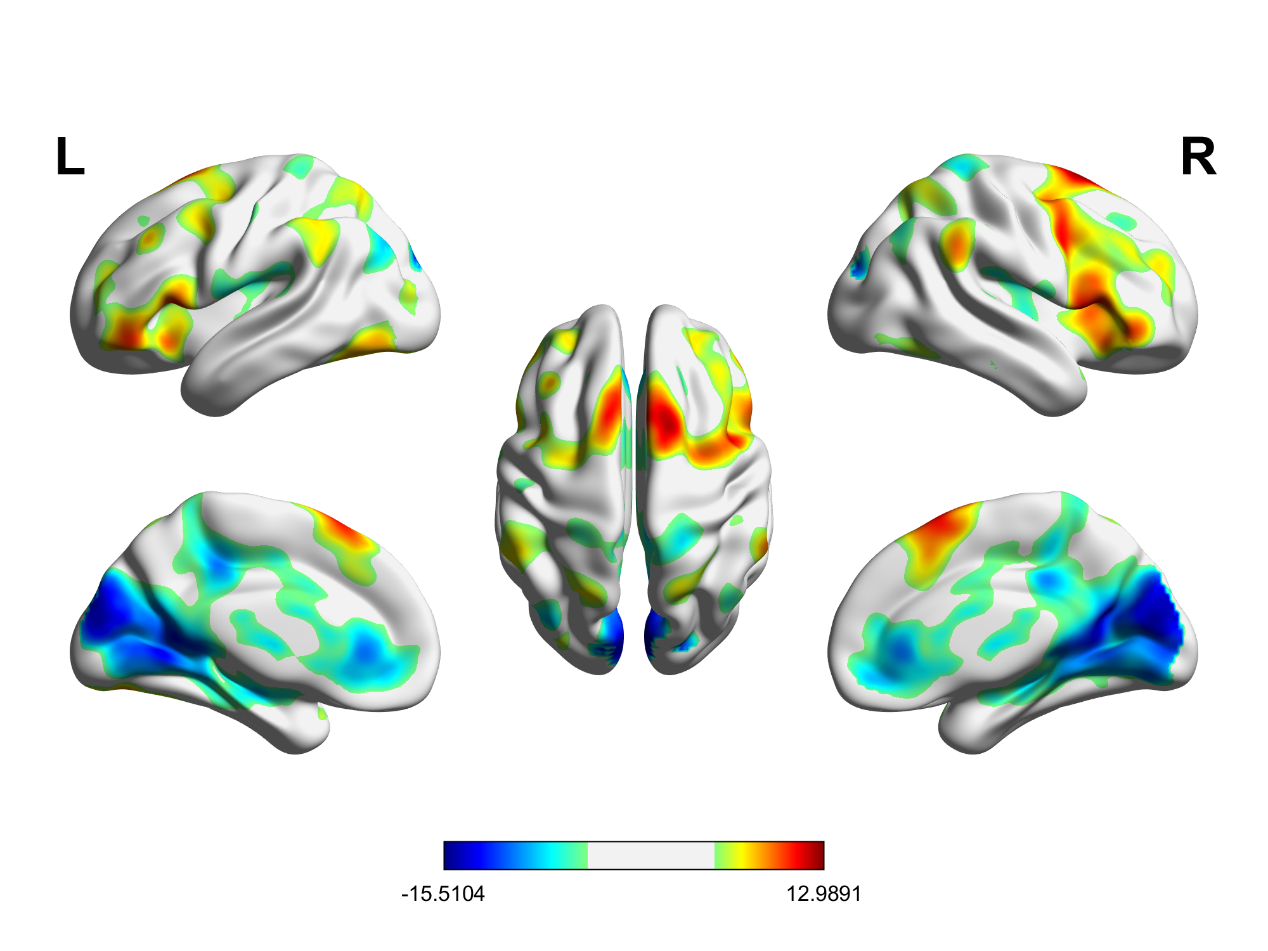

**Fig. 1.** **Regions showing a difference in activity between No-Think and Think trials** (No-Think > Think). The unit of the color map bar is the T value. Statistical parametric maps were thresholded at *P_FWE_* < .05 and rendered using the BrainNet Viewer package.
